# Cingulate prediction of response to antidepressant and cognitive behavioral therapies for depression: Theory, meta-analysis, and empirical application

**DOI:** 10.1101/2020.12.02.407841

**Authors:** Marlene V. Strege, Greg J. Siegle, Kymberly Young

**Affiliations:** Virginia Polytechnic Institute and State University; University of Pittsburgh Medical Center; University of Pittsburgh School of Medicine

## Abstract

**Objective:** In the interest of precision medicine, we sought to derive preclinical markers of neural mechanisms associated with treatment response in unipolar depression, separated by treatment type.

**Methods:** We conducted separate neuroimaging meta-analyses of neural predictors for response to Cognitive Behavioral Therapy (CBT) and Selective Serotonin Reuptake Inhibitors (SSRIs). We assessed whether reactivity of derived regions predicted clinical change in a preference trial of patients with major depressive disorder (MDD) who received CBT (n = 61) or SSRIs (n = 19).

**Results:** The meta-analyses yielded regions within the perigenual (pgACC) and subgenual anterior cingulate cortex (sgACC) associated with SSRI and CBT response, respectively. In our sample, reactivity of the sgACC region was prognostic for response to CBT, but neither cingulate region was prognostic for response to SSRIs using a linguistic task; most prognostic SSRI studies used images. An exploratory analysis revealed a pgACC region for which reactivity to images was prognostic for response to SSRIs.

**Conclusions:** Results suggest that neural reactivity of the sgACC and pgACC are associated with CBT and SSRI response for unipolar depression. Further research incorporating methodological considerations is necessary for translation.

## Introduction

Precision medicine is increasingly central to healthcare, though is far from the standard of care in psychiatry where pre-treatment biological assessments are rarely used. Neuroimaging has the potential to identify mechanisms that could guide development of clinical measures and targeted interventions. This use is hindered by variable results, data collection, and analytic methods. Meta-analyses (1,2) could address these issues. To obtain possibly predictive indicators, we subjected published neuroprediction studies to separate meta-analyses for two common interventions for depression - Cognitive Behavioral Therapy (CBT) and Selective Serotonin Reuptake Inhibitors (SSRIs). We further examined whether obtained results applied to a preference neuroimaging CBT/SSRI trial for major depressive disorder (MDD) (“confirmatory sample”). We focus on the perigenual and subgenual regions of the anterior cingulate cortex (pgACC and sgACC), which have been implicated in treatment outcome in depression (for metaanalyses see (1) and (2)).

Our first question (Q1), examined via our meta-analysis, is whether these regions are differentially predictive in the literature. We have, based on the state of the literature a decade ago, hypothesized (3) that lower pre-treatment sgACC reactivity to negative information is associated with greater symptom improvement (4,5) and that greater pre-treatment pgACC and sgACC reactivity to negative emotional stimuli are implicated in better antidepressant medication outcomes (6–9). Our second question (Q2) regards the generalizability of these results, which we considered by examining the predictive utility of the derived regions in the confirmatory sample. If robust regions are detected meta-analytically which strongly predict in the confirmatory sample (Q2a) we could suggest the literature has identified preclinical indicators that are useful at the single subject level. If regions are detected that require more work to replicate in the confirmatory sample (Q2b), we can suggest the literature has identified mechanistic targets for understanding vulnerability to treatment failure and potentially for pretreatment remediation but is not yet ready for these regions to be used as clinical predictors in new studies.

## Method

### Meta-Analytic Methodology

To obtain studies for the meta-analyses, we reviewed all articles included in recent reviews of the neuroimaging depression treatment outcome prediction literature through March 2019 (1,2,10,11). We also conducted supplemental literature searches via PubMed and Google Scholar using relevant terms such as “depression,” “neural predictors,” “treatment,” and “cognitive behavioral therapy.” Articles were included if they reported pre-treatment neural activation (fMRI or PET) as prognostic or predictive of treatment outcome in depression. We limited the meta-analyses to studies that assessed treatment response to SSRIs or CBT (including variants such as behavioral activation), provided coordinates for their region results, and were published before March of 2019. We also limited the meta-analyses to studies with emotional stimuli that reported on activation (not exclusively connectivity) consistent with our theoretical framework. Table 1 lists included articles; Supplement Table 1 lists obtained articles that were excluded and reasons for exclusion.

**Table 1.** List of papers included in meta-analyses

| Treatment Type | Study | Sample (N) |
| --- | --- | --- |
| CBT and Related |  |  |
|  | Costafreda et al., 2009 (31) | MDD adults (16) |
|  | Dichter et al., 2010 (32) | MDD adults (12)<br>Controls (15) |
|  | Doerig et al., 2016 (33) | MDD adults (21) |
|  | Fu et al., 2008 (34) | MDD adults (16) |
|  | Ritchey et al., 2011 (35) | MDD adults (22)<br>Controls (14) |
|  | Siegle et al., 2006 (4) | MDD adults (14) |
|  | Siegle et al., 2012 (5) | MDD adults (49) |
|  | Straub et al., 2015 (36) | MDD adolescents (22) |
|  | Yoshimura et al., 2014 (37) | MDD adults (23) |
| SSRI |  |  |
|  | Chen et al., 2007 (6) | MDD adults (17) |
|  | Godlewska et al., 2018 (7) | MDD adults (32) |
|  | Miller et al., 2013 (21) | MDD adults (15) |
|  | Roy et al., 2010 (9) | MDD adults (20) |
|  | Ruhe et al., 2012 (38) | MDD adults (20) |
|  | Spies et al., 2017 (39) | MDD adults (22) |
|  | Victor et al., 2013 (8) | MDD adults (10) |
| CBT & SSRI |  |  |
|  | Forbes et al., 2010 (40) | MDD adolescents (6) |

From included articles we subjected all coordinates reported as predictive of treatment response in depression to GingerALE (version 3.0.2), a software package that conducts activation likelihood estimation (ALE) meta-analyses using coordinate-based data. We conducted separate meta-analyses for CBT and SSRI studies. If a study reported a region that was predictive of response to both treatments, coordinates were included in both meta-analyses.

For both meta-analyses, we set cluster-level familywise error to 0.05, voxelwise p-value threshold to 0.005. This is consistent with prior work (12) suggesting that a more conservative threshold (e.g., voxel threshold of p < .001) may be too conservative for neuroimaging studies in clinical samples. Threshold permutations were set to 1,000, with GingerALE’s dilated 2mm mask size parameter.

### Confirmatory sample

Our confirmatory sample consisted of 98 individuals with MDD recruited in an expanded sample from our prior fMRI CBT-outcome-prediction study (5) and a parallel preference-based SSRI-outcome-prediction study (not yet published) (Consort Diagram in Supplement 2). The sample for the CBT study was largely that used in one of the studies in the meta-analysis (5) but was slightly enlarged as additional data were collected after the initial article submission and missing clinical response variables were imputed. Participants with MDD were thus recruited from two clinical trials (ClinicalTrials.gov: NCT00183664; NCT00787501) and were treated with either SSRIs prescribed by a psychiatrist or CBT by 6 community clinicians (Ph.D.’s, M.D.’s, M. Ed.’s, LCSW’s) who ranged widely in CBT experience (described fully in (5)). Participants described no health problems, eye problems, or psychoactive drug abuse in the past six months and no history of psychosis, manic, or hypomanic episodes. Participants had not used antidepressants within two weeks of initial testing (six weeks for fluoxetine) due to either medication naivety or supervised withdrawal from unsuccessful medications. Participants reported no excessive use of alcohol in the two weeks prior to testing and scored in the normal range on a cognitive screen (13), VIQ-equivalent > 85. Of the initial 98 patients, 80 individuals completed treatment and are reported on in this paper.

### Protocol and Treatment Procedures

Full details on treatment procedures are provided in Supplement 3. Briefly, after obtaining written informed consent, participants completed a diagnostic interview (Structured Clinical Interview for DSM-IV, SCID) (14), and fMRI assessment as in (5). The first N=17 depressed participants were part of a CBT trial (15). Subsequent participants selected CBT (N=57) or medication (N=24) trial by preference. Of the combined sample, CBT (N=61) and SSRI (N=19) completed treatment. We conducted analyses both in a sample restricted to completers as well as a sample including non-completers (Supplements 7, 8), consistent with previous work (5). CBT and medication were given at the same frequency; 2 sessions/week for the first four weeks followed by 8 weekly sessions for “early-responders” (HRSD reduction <40% at session 9; 16 total sessions) or 2 sessions/week for the first 8 weeks followed by 4 weekly sessions for non-early-responders (20 total sessions). CBT followed Beck’s (16) guidelines (5). Pharmacotherapy sessions involved only general mood check, Hamilton Rating Scale for Depression (HDRS) (17) assessment, psychoeducation on drug effects, side effects assessments, and treatment regime change discussion. The default prescribed medication was escitalopram (N=15; 10-30mg/day), or if that had previously been failed, then a comparable dose of sertraline (N=4), fluoxetine (N=4), or, in N=1 case, another SSRI/SNRI. Participants were scanned again within 2 weeks of completion.

### Clinical response

Our primary outcome measure was the Beck Depression Inventory (BDI-II) (18), a widely-employed 21-item self-report measure of depression, assessing symptom severity on a 0 (not present) to 3 (severe) scale with strong psychometric properties (19,20). Consistent with prior work (5), we considered improvement as both BDI-II change (post-pre) as well as residual severity, calculated as final severity controlling for initial severity. Six participants were missing pre-treatment BDI scores, and 13 participants were missing post-treatment BDI scores. Imputation procedures are described in Supplement 4.

### fMRI task

#### Apparatus

Twenty-nine 3.2mm slices were acquired parallel to the AC-PC line (3T Siemens Trio, T2*-weighted images depicting BOLD contrast; posterior-to-anterior, TR=1500ms, TE=27ms, FOV=24cm, flip=80), yielding 8 whole-brain images per 12 second trial. Stimuli were displayed in black on a white background via a back-projection screen (.88° visual angle). Responses were recorded using a Psychology Software Tools™ glove.

#### Personal relevance rating task (PRRT)

As described fully in our previous publication on this sample (5), in 60 slow-event related trials, participants viewed a fixation cue (1 s; row of X’s with prongs around the center X) followed by a positive, negative, or neutral word (200 ms; only negative words analyzed here), followed by a mask (row of X’s; 10.8 s). Participants pushed a button for whether the word was relevant, somewhat relevant, or not relevant to them or their lives (button orders balanced across participants), as quickly and accurately as they could. Participant-generated and normed words were used as in our previous studies of depression (4). For fMRI data preparation see Supplement 4.

### Analyses

Analyses involved examination of whether Q2a) activity averaged over the meta-analytically derived regions were predictive in the directions suggested by the meta-analysis in the new dataset (i.e., whether the meta-analytic regions could serve as pre-clinical markers), Q2b) whether subregions or regions close to the meta-analytic regions were predictive (i.e., whether meta-analytic regions could guide mechanistic investigations useful for treatment refinement in new studies).

## Results

### Q1. Meta-analytic identification of brain regions prognostic for treatment outcome in depression

The ALE meta-analysis for CBT and related treatments included 10 studies and resulted in 2 significant clusters (Figure 1a). One cluster in the right sgACC (size = 4158.65 mm^3^, (MNI centroid coordinates are used throughout this manuscript) *x* = 7, *y* = 23, *z* = −12), which involved activation shared by 3 studies. After applying a gray matter mask, this became a 2488.96 mm^3^ cluster with the same centroid coordinates, see Figure 1b. The other cluster was in the right amygdala (3548.97 mm^3^, *x* = 22, *y* = −1, *z* = −22) and corresponded to shared activation of 4 studies. The meta-analysis for SSRI treatment studies included 8 articles and resulted in 2 significant clusters (Figure 2a) including the right pgACC (8745.13 mm^3^, *x* = 19, *y* = 36 *z* = 0) and right caudate (3670.22 mm^3^, *x* = 20, *y* = 12, *z* = 19). After applying a gray matter mask to the pgACC-centered cluster, we applied a minimum cluster threshold of 20 voxels (this is in addition to the aforementioned cluster corrections and was set to limit reporting on small clusters created from applying the gray matter mask). This yielded 5 clusters spanning the right pgACC, (746.51 mm^3^, *x* = 15, *y* = 41, *z* = 0, see Figure 2b) along with activations in the right caudate and other medial frontal regions (non-ACC clusters in Supplement 5).

**Figure 1.**
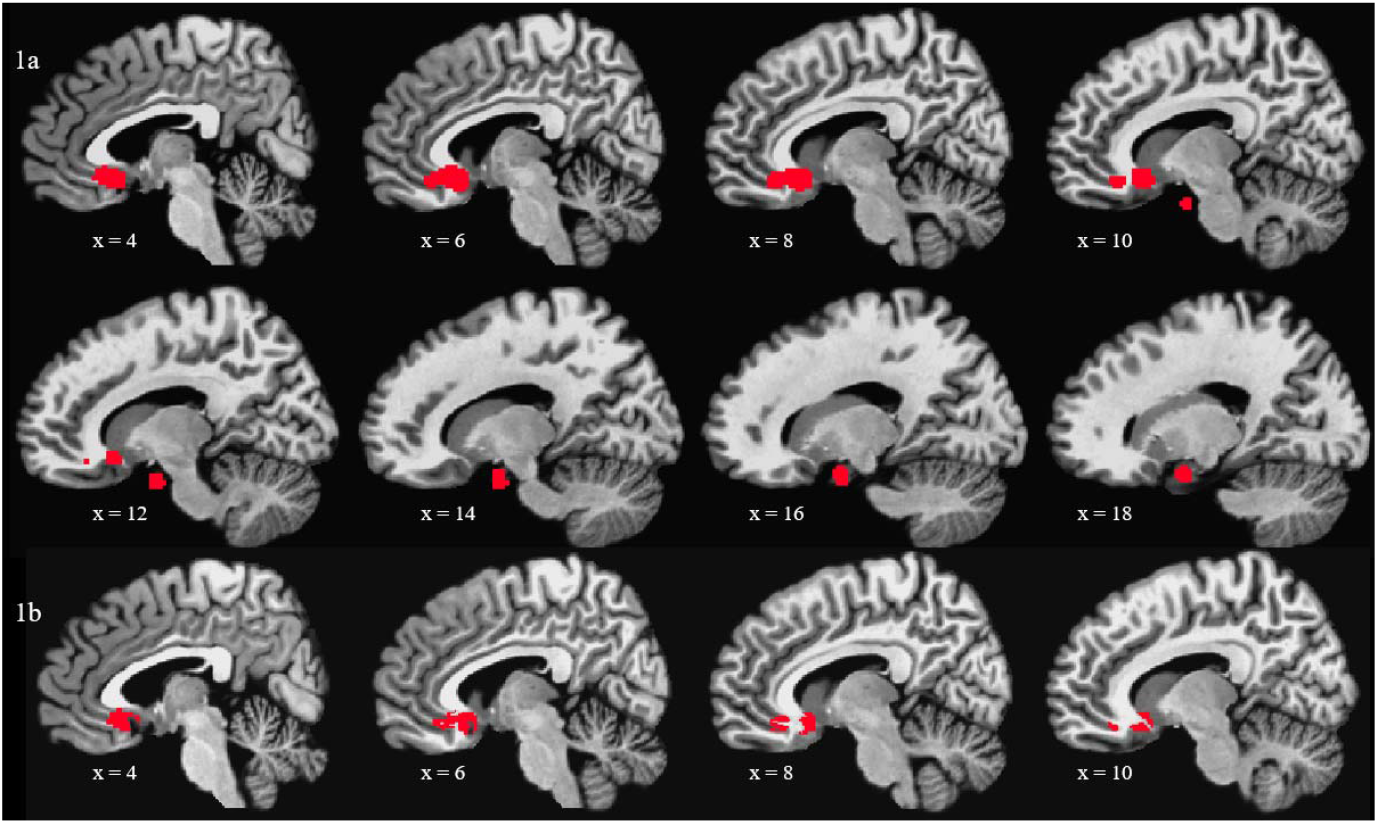
ALE meta-analysis results from CBT studies in depression Notes: a. Significant clusters from the ALE meta-analysis of CBT studies in depression b. Resulting subgenual cingulate region after graymatter mask of ALE meta-analysis significant clusters

**Figure 2.**
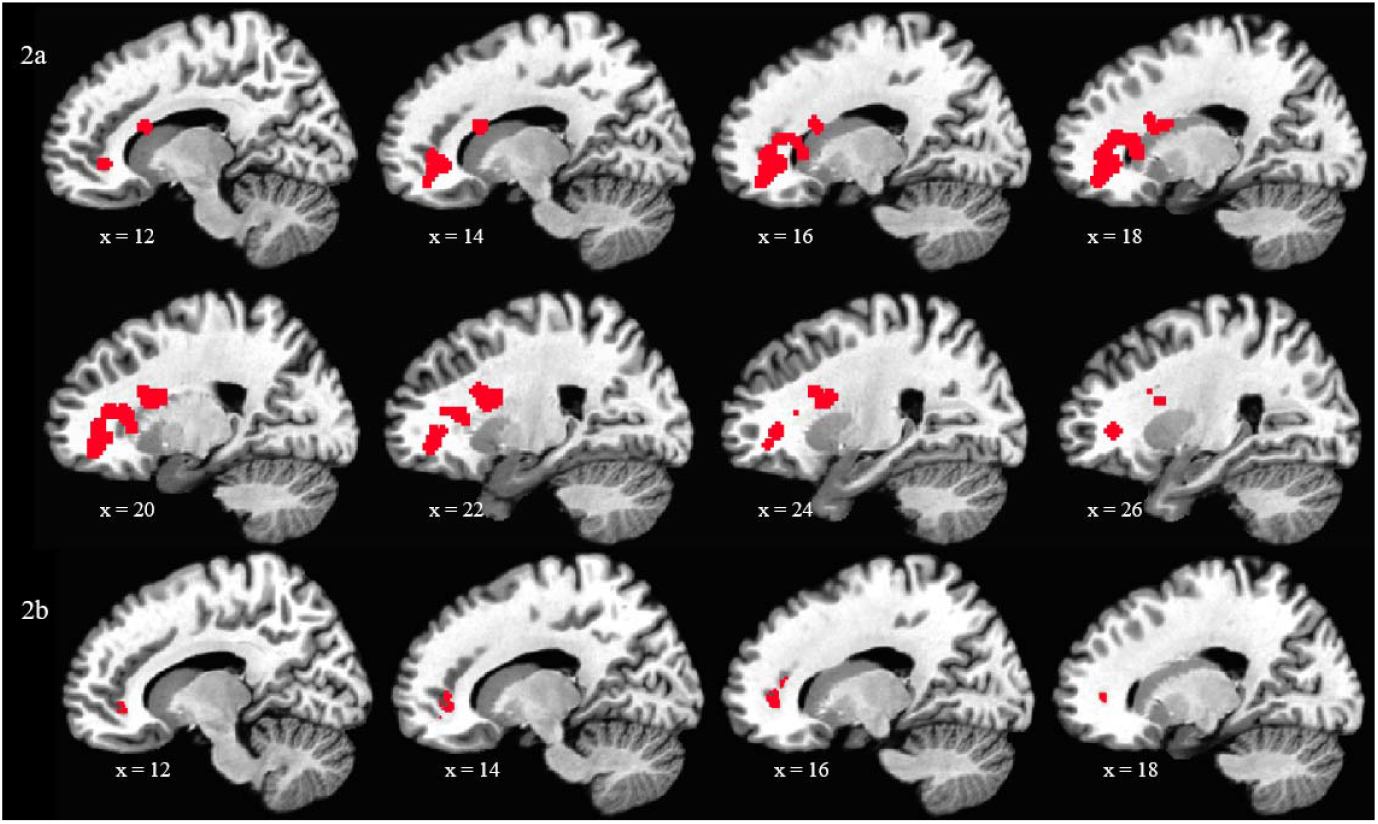
ALE meta-analysis a) results of SSRI studies in depression and b) perigenual cingulate region surviving gray matter mask Notes: a. Significant clusters from the ALE meta-analysis of SSRI studies in depression b. Resulting perigenual cingulate region after graymatter mask of ALE meta-analysis significant clusters

### Q2. Prospective application of meta-analytically derived regions to a new clinical dataset

#### Demographics of the clinical sample

Treatment groups did not differ on gender, age, ethnicity, number of depressive episodes, or depressive symptoms (BDI-II). The CBT patient sample reported more years of education than the SSRI sample, *t*(76) = 3.03, *p* = .003, Hedges’ g = 0.79 (Table 2). Similar demographics for the entire sample (not limited to those who completed treatment) are reported in Supplement 6.

**Table 2.** Demographics of unipolar depression patients, limited to patients who completed the protocol

|  | **Whole Patient Sample (N = 80)** | **CT Patient Sample (N = 61)** | **SSRI Patient Sample (N = 19)** |
| --- | --- | --- | --- |
| Gender | Female (N = 56, 70.00%) | Female (N = 43, 70.49%) | Female (N = 13, 68.42%) |
| Age | 36.66 (10.69) years | 36.23 (10.23) years | 38.05 (12.24) years |
| Ethnicity | Caucasian (N = 64, 80.00%)<br>Non-Caucasian (N = 16, 20.00%) | Caucasian (N = 49, 80.33%)<br>Non-Caucasian (N = 12, 19.67%) | Caucasian (N = 15, 78.95%)<br>Non-Caucasian (N = 4, 21.05%) |
| Education | 15.38 (2.32) years | 15.81 (2.30) years* | 14.05 (1.87) years* |
| Depressive episodes | 3.64 (2.69) | 3.70 (2.74) | 3.44 (2.62) |
| Beck Depression Inventory - II (BDI-II) | Pre Treatment: 29.69 (8.58)<br>Post Treatment: 11.68 (9.52) | Pre Treatment: 30.15 (8.29)<br>Post Treatment: 11.80 (9.37) | Pre Treatment: 28.20 (9.55)<br>Post Treatment: 11.32 (10.24) |

Note. Means for age, education, number of depressive episodes, and Hamilton Depression Rating Scale are followed by standard deviations in parentheses. \* Denotes that the CT and SSRI patient samples statistically differ on this variable. CT sample education (N=59), number of depressive episodes (N=56). SSRI sample number of depressive episodes (N=18).

### Q2a. Prediction based on activity averaged over meta-analytically derived ROIs

#### sgACC ROI from CBTstudies

For patients who received CBT, sgACC ROI responses to linguistic stimuli were weakly but significantly prognostic for BDI change scores *R*^2^ = .07, *F*(1,59) = 4.33, *p* = 0.042 (Figure 3). Less sgACC reactivity to negative stimuli was associated with stronger response to CBT. Although comparable in effect size, this relationship failed to meet statistical significance for BDI residuals (*p* = .057). This ROI was not prognostic for BDI change scores *R*^2^ < .01, *F*(1,18) = <.01, *p* = 0.989 or residuals *R*^2^ < .01, *F*(1,18) = <.01, *p* = 0.989 in the SSRI sample.

**Figure 3.**
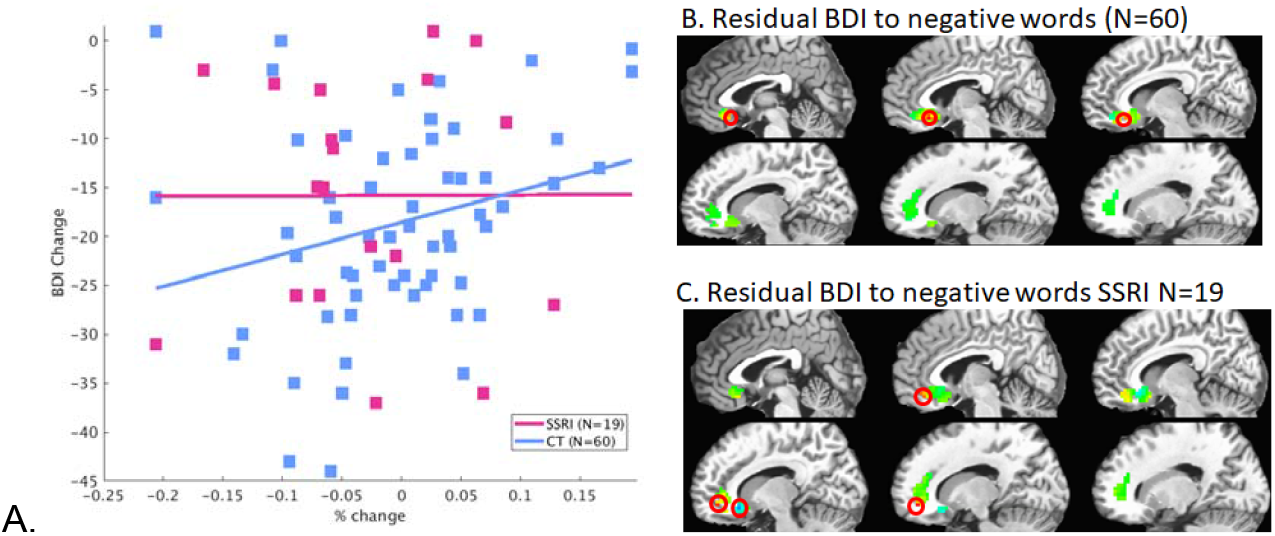
A) Therapy ROI (subgenual cingulate) and BDI-II change scores for those who completed treatment. B,C) Associations (R^2) of residual BDI with voxelwise reactivity (scans 2-7) in response to negative words (as in (5)) within pgACC and sgACC regions with residual sympomatology. Thus red values indicate associations of higher activity with less negative residual sympatomatology (poorer recovery); lower activity is associated with better recovery. Significant (p<.05) associations are circled. In the CBT group, lower sgACC activity to words was associated with improved recovery. In the SSRI group, higher sgACC activity and lower pgACC activity to words was associated with improved recovery. For both groups R^2s as high as 0.36 were observed in some voxels.

#### pgACC ROI from SSRIstudies

pgACC ROI responses to linguistic stimuli were not significantly prognostic for BDI change scores *R*^2^ = 0.08, *F*(1,18) = 1.52, *p* = 0.234 or residuals *R*^2^ = 0.15, *F*(1,18) = 2.87, *p* = 0.108, though effect sizes were comparable to or larger than the sgACC effect for CBT responders. In contrast to the meta-analytic association, higher reactivity was associated with higher residual symptomatology. The ROI also was not associated with greater treatment response for CBT patients (change scores: *R*^2^ < .01, *F*(1,60) = 0.15, *p* = 0.705, residuals: *R*^2^ < .01, *F*(1,60) = 0.04, *p* = 0.842). Similar effects (Supplements 7, 8) were found when examining the entire clinical dataset (not limited to those who completed treatment).

### Q2b. Prediction based on indices derived from the meta-analytic regions

#### Subregions

Our first consideration involved whether subregions of the meta-analytic region were more strongly predictive than the overall region. As shown in Figure 3B,C, subregions of each of the meta-analytically derived regions were more strongly predictive (R^2’s up to 0.36) than the average over the regions, with some parts of these regions being negatively and other parts being positively associated with response to each treatment modality. The largest part of the sgACC was associated with outcome for CBT in the meta-analytically predicted direction (less activity was associated with better response). Part of the sgACC was associated, in the opposite direction, with outcome for SSRIs, but unlike the meta-analysis, the only significant associations within the pgACC were the same as for CBT.

#### Potential moderation of SSRI prediction by stimulus modality

The lack of correspondence of the current sample SSRI prediction with the meta-analysis prompted us to consider what was different between this sample and the literature. Of the SSRI studies that contributed to the perigenual cluster (6–9,21), all except (21) found that greater reactivity of the pgACC was prognostic of better treatment outcome, and all except (21) used picture stimuli. In contrast, (21) used linguistic stimuli similar to our study and found that less reactivity to negative words predicted better treatment response. As our SSRI sample also completed Hariri’s emotional faces task (22), we were able to test whether stimuli modality moderated prediction. In that task, increased pgACC reactivity to fearful and angry faces was prognostic for decreased depressive severity as in the meta-analysis in the region as a whole (Supplement 9).

#### Public dissemination of results

So that other researchers can examine our meta-analytically derived regions on their own data, we have made them publicly available at: https://www.pitt.edu/~gsiegle/strege_metaanalysis_rois.tar.gz.

## Discussion

Our primary question was whether meta-analytically derived cingulate regions were differentially predictive of response to SSRIs and CBT. Indeed, our meta-analysis revealed that anterior cingulate subregions were broadly prognostic for stronger response to treatment for depression, which is largely consistent with other meta-analyses (1,2). CBT outcome for major depressive disorder was associated with decreased right sgACC reactivity, whereas SSRI outcome was associated with increased right pgACC as well as caudate reactivity. Our second question (Q2) regarded the robustness of these regions when applied to a single study which used a linguistic task. Regarding the meta-analytic regions considered as a whole, in the simplest analyses (Q2a), the CBT-study derived sgACC ROI reactivity to negative words was moderately associated with clinical response to CBT in the single study - not strongly enough to be considered a robust indicator for use in clinics (R^2’s < .3). The SSRI-study derived pgACC ROI reactivity to negative words was not associated with clinical response to SSRIs in that sample. That said, we also considered whether slightly more nuanced analyses would reveal interesting mechanistic generalizability (Q2b). This analysis revealed 1) that subregions of the meta-analytically derived regions were differentially predictive for the new sample and 2) that the apparent differentiation examined in the meta-analysis may be due to the primary use of linguistic stimuli in CBT studies and images in SSRI studies. Indeed, pgACC reactivity to negative images was associated with clinical response (Supplement 9). We also found the right caudate was associated with response to SSRIs. The caudate’s roles in reward reactivity (23) and depression (24) could help to explain this finding.

That the meta-analytic finding for pgACC prediction of SSRI response was supported by pictures, and not word stimuli in our local sample could suggest that differential meta-analytic results for treatment modality are actually due to systematic differences in stimulus type employed by the literature. Indeed, all but one (21) of the SSRI studies contributing to the meta-analytic pgACC cluster used picture stimuli (6–9). The study that used linguistic stimuli (21) found the opposite effect direction, consistent with our study. Results in our sample were consistent with this explanation though our SSRI sample was atypical in other potentially important ways, e.g., having weekly provider meetings. Further work examining neuroprediction associations with stimulus modality and treatment session frequency would help clarify the influence of study methodology on prognostic findings.

Results suggest that the meta-analytically derived regions need refinement before being translated to the clinic. Extension to robust predictors that work in novel datasets, and understanding of methodological moderation effects, pre-treatment scans or associated physiological, EEG (25), or self-report measures could help to determine the intervention patients should be encouraged to receive, particularly if they are amenable to both therapy and medications.

An avenue for future research more strongly supported by the current investigation involves the potential to target the proposed regions to optimize likely treatment response based on an individual’s treatment preference. For example, (26) found that healthy individuals could successfully downregulate reactivity to negative pictures within sgACC with neurofeedback training and the strategy of increasing positive mood. Other studies have trained both healthy participants and patients with schizophrenia to increase the activity in the sgACC with neurofeedback while recalling positive memories and imagining playing sports and listening to music (27,28). Thus, perhaps, treatment-seeking individuals could benefit from fMRI-guided pre-treatment neurofeedback of anterior cingulate subregion reactivity (to increase or decrease reactivity with respect to the level found in healthy individuals). While emerging evidence suggests fMRI neurofeedback may be an effective intervention for depression (29), one common critique is that it is too expensive to ever be widely clinically implemented (30); however for patients who are not predicted to respond to any intervention, fMRI neurofeedback may be a particularly viable approach.

Results should be interpreted with consideration of multiple limitations. The metaanalysis had few studies of CBT including two from our lab, one of which was on a subset of participants in our confirmatory sample, making it a non-independent replication. Nearly all of the included studies had small sample sizes. Our confirmatory sample had non-random treatment assignment; it was a convenience sample of some participants coming from a preference trial and others from a CBT-only trial. As we did not assess other depression interventions, if patients were predicted to not respond to either CBT or SSRIs, our results do not indicate whether they would respond to other interventions, such as electroconvulsive therapy or whether there are pretreatment remediations that could be employed, such as neurofeedback, to change their predictive status.

These limitations notwithstanding, the current data suggest that neuroimaging provides promising neural correlates (sgACC and pgACC) of treatment response to two of the most common and well-validated interventions for unipolar depression, CBT and SSRIs. They are not yet robust as predictors for new studies without attending to subregions and stimulus type, but can serve to guide intervention development and pre-treatments. Attending to stimulus type and treatment session frequency could make it more likely our meta-analytically derived regions generalize in future studies.

