## Supplement for "Cingulate prediction of response to antidepressant and cognitive behavioral therapies for depression: Theory, meta-analysis, and empirical application"

Contents:

Supplement 1. Excluded articles and reasons for exclusion	**2**

Supplement 2. CONSORT Diagram	**6**

Supplement 3. Protocol and Treatment Procedures - all the details	**6**

Supplement 4. Analysis details	**8**

A. Missing data imputation	8

B. fMRI data preparation	8

Supplement 5. Non-ACC clusters surviving graymatter masking of pgACC-centered SSRI ROI, centroid MNI coordinates reported	**9**

Supplement 6. Demographics of MDD patients, entire sample	**10**

Supplement 7. Application of meta-analytically derived regions to entire clinical dataset (not limited to those who completed treatment)	**11**

Supplement 8. Therapy ROI (subgenual cingulate) and BDI change scores for entire sample	**12**

Supplement 9. Prognostic accuracy of neural responses to faces for SSRI response in the empirical sample	**13**

References to the Supplement	**16**

### Supplement 1. Excluded articles and reasons for exclusion

**Structural Predictors**

Medication Papers

(Baldwin et al., 2004)

(Frodl et al., 2004)

(Frodl et al., 2008)

(Gong et al., 2011)

(Gunning-Dixon et al., 2010)

(Hoogenboom et al., 2014)

(Hsieh et al., 2002)

(Iosifescu et al., 2006)

(Jung et al., 2014)

(Khalaf et al., 2015)

(Li et al., 2010)

(Liu et al., 2012)

(MacQueen et al., 2008)

(Patankar et al., 2007)

(Phillips et al., 2012)

(Vakili et al., 2000)

Therapy Papers

(Costafreda et al., 2009)

(Mackin et al., 2013)

**Connectivity-Based Predictors**

Medication Papers

(Alexopoulos et al., 2012)

(Andreescu et al., 2013)

(He et al., 2016)

(Lisiecka et al., 2011)

(L.-J. Wang et al., 2014)

(L. Wang et al., 2015)

Therapy Papers

(Dunlop et al., 2017)

(Straub et al., 2017)

**Change-Based Correlates**

Medication Papers

(Cullen et al., 2016)

(Godlewska et al., 2016)

(Mayberg et al., 2000)

Therapy Papers

(Goldapple et al., 2004)

Combined Treatment Papers

(Kennedy et al., 2007)

**Alternative Non-Neuroimaging Methods**

Medication Papers

(Korb et al., 2009)

(Mulert et al., 2007)

(Pizzagalli et al., 2001)

(Yeh et al., 2015)

Therapy Papers

(Amsterdam et al., 2013)

(Sanacora et al., 2006)

(Tiger et al., 2014)

Combined Treatment Papers

(Hirvonen et al., 2011)

**Non-SSRI Medication**

(Brannan et al., 2000)

(Crane et al., 2017)

(Davidson et al., 2003)

(Delaveau et al., 2016)

(Frodl et al., 2011)

(Fu et al., 2015)

(Furey et al., 2013)

(Furey et al., 2015)

(Keedwell et al., 2010)

(Konarski et al., 2009)

(Little et al., 2005)

(López-Solà et al., 2010)

(Rizvi et al., 2013)

(Salvadore et al., 2009)

(Samson et al., 2011)

(Szczepanik et al., 2016)

(Toki et al., 2014)

(Williams et al., 2015)

**Non-CBT Therapy**

(Roffman et al., 2014)

**Alternative Treatment**

(Hernández-Ribas et al., 2013)

**Coordinates Not Provided**

(Brody et al., 1999)

(Carl et al., 2016)

**Task without Emotional Stimuli**

(Gyurak et al., 2016)

(Langenecker et al., 2007)

(Marquand et al., 2008)

(McGrath et al., 2013)

(McGrath et al., 2014)

(Milak et al., 2009)

(Saxena et al., 2003)

(Wagner et al., 2010)

(Walsh et al., 2007)

(Thompson et al., 2015)

### Supplement 2. CONSORT Diagram

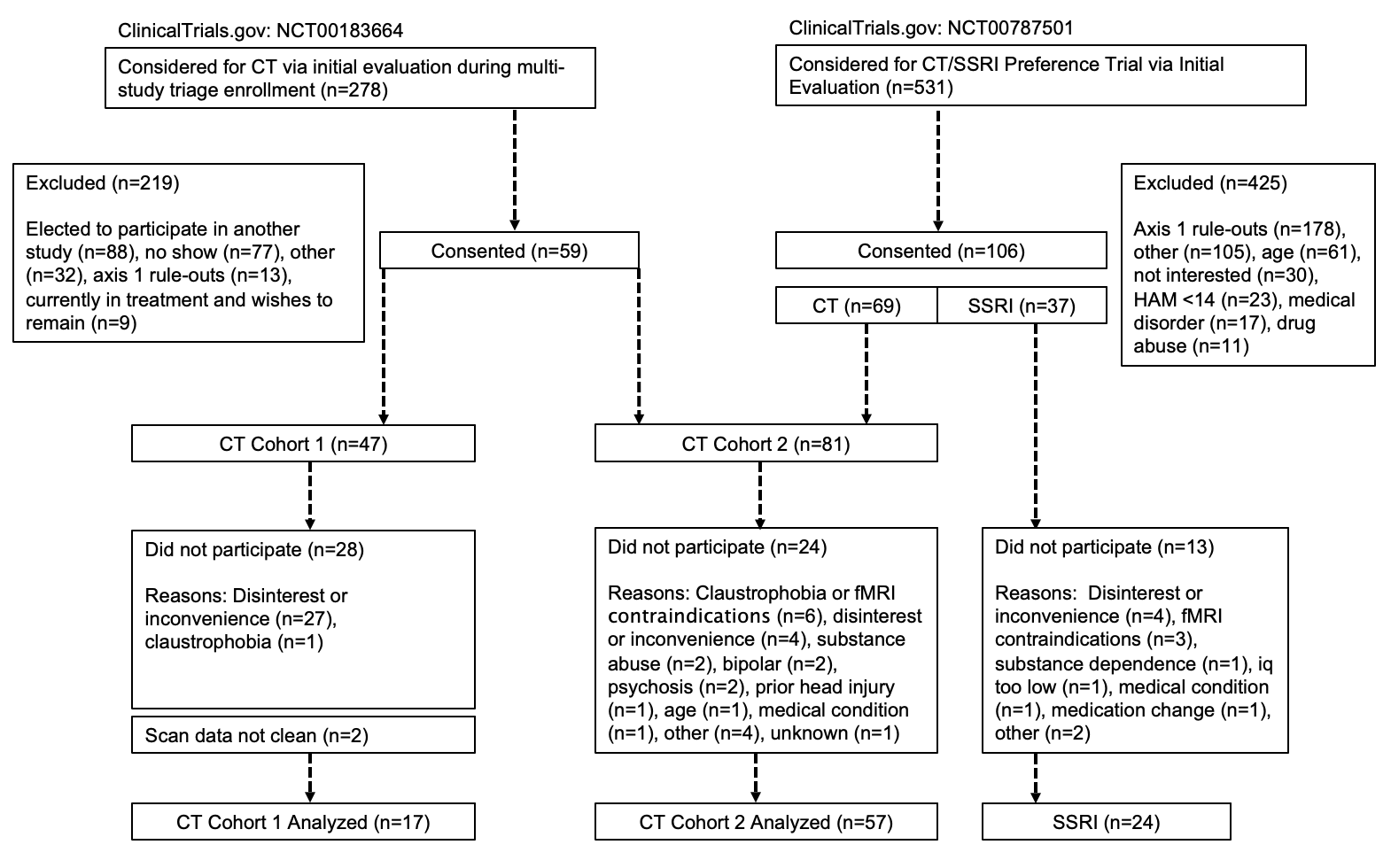

### Supplement 3. Protocol and Treatment Procedures - all the details

After describing the study, we obtained IRB-approved written informed consent. We then conducted a diagnostic interview (Structured Clinical Interview for DSM-IV, SCID) (First et al., 1995), vision test, and unrelated physiological assessment. On a separate day, participants underwent a battery of fMRI tasks administered in counterbalanced order fully described in (Siegle et al., 2012). Depressed participants then received CBT or medication. The first N=17 participants were part of a CBT trial (Jarrett et al., 2013). Subsequent participants selected CBT (N=57) or medication (N=24) trial by preference. CBT and medication were given at the same visit frequency; 2 sessions/week for the first four weeks followed by 8 weekly sessions for “early-responders” (HRSD reduction <40% at session 9; 16 total sessions) or 2 sessions/week for the first 8 weeks followed by 4 weekly sessions for non-early-responders (20 total sessions). CBT followed Beck’s (Beck et al., 1979) guidelines as described in (Siegle et al., 2012). Pharmacotherapy sessions were 30-45 minutes in length and were conducted by a psychiatric nurse who inquired about general mood status, did a Hamilton Rating Scale for Depression (HDRS) (Hamilton, 1960) assessment, and provided psychoeducation about medication effects and adverse effects. A psychiatrist consulted with the nurse and patient for the final 5-10 minutes of the session. Symptoms, adverse effects, and treatment progress were reviewed with the psychiatrist and treatment recommendations were made. No psychotherapy or supportive counseling was provided. The discussion of subjects’ thoughts, feelings, and/or behaviors by the pharmacotherapy team was forbidden. The default prescribed medication was escitalopram (N=15; 10mg/day at start, increased to 30mg/day by week 6 if no response, max dose M=19.33mg, SD=4.58mg), or if that had previously been failed, then a comparable dose of sertraline (N=4, max dose M=125.00mg, SD=28.87mg) or fluoxetine (N=4, max dose M=27.50mg, SD=9.57mg). One participant changed medications during the trial from escitalopram to fluoxetine (counted as part of the fluoxetine group above), and we do not have medication information for one individual. Within two weeks of completion (week 16 for controls) all participants completed the same fMRI protocol and treatment outcome measures again.

### Supplement 4. Analysis details

#### A. Missing data imputation

For the clinical sample for the primary analyses, BDI scores were missing for 6 patients at the pre and 13 patients at the post treatment assessment. Of these, 6 pre BDI scores and 11 post BDI scores are missing due to licensing issues that resulted in the BDI not being able to be collected in the interim. Reasons that the remaining 2 post BDI scores were missing were not noted. For imputation of missing data, we submitted related measures including the HDRS (Hamilton, 1960), State-Trait Anxiety Inventory (Spielberger, 2010), the General Distress Depressive Symptoms and Anhedonic Depression subscales of the Mood and Anxiety Symptom Questionnaire (Watson et al., 1995), and the Rumination subscale of the Response Styles Questionnaire (Nolen-Hoeksema, 1991) to the SPSS multiple imputation procedure (averaging across 5 imputations).

#### B. fMRI data preparation

We followed standard preprocessing described fully in (Siegle et al., 2012) (slice time correction, motion correction, linear detrending, voxelwise outlier rescaling, conversion to percent-change, temporal smoothing (5 point middle peaked filter), 32 parameter nonlinear warping the Montreal Neurological Institute Colin-27 brain, and spatial smoothing (6mm FWHM), response time-series variability normalization across scanners). We computed peak and sustained responses to negative words as the mean of the 4^th^-7^th^ images (henceforth “scans”) of each negative-word trial minus the trial’s first (pre-stimulus) scan acquired while the fixation cue was on the screen, i.e., 6-10.5 seconds after stimulus onset, or 5-9.5 seconds following the onset of the word, consistent with our previous work (Siegle et al., 2006, 2012).

To assess neural reactivity outliers, we conducted box-and-whisker plots and examined each value’s position relative to the interquartile range (IQR). This resulted in the removal of two data points from the regressions using the CBT-derived region, as these data points were > 3 x IQR and appeared separate from the distribution upon visual inspection of the box-and-whisker plot. Additional reactivity data points outside of 1.5 x IQR were winsorized, rescaled to the maximum value, same direction.

### Supplement 5. Non-ACC clusters surviving graymatter masking of pgACC-centered SSRI ROI, centroid MNI coordinates reported

| Location | Size (mm^3^) | x | y | z |
| --- | --- | --- | --- | --- |
| Right Caudate | | | | |
|  | 815.80 | 18 | 23 | 4 |
|  | 670.30 | 18 | 10 | 16 |
| Right Middle Frontal Gyrus | | | | |
|  | 90.07 | 20 | 45 | -14 |
| Right Medial Frontal Gyrus |  |  |  |  |
|  | 41.57 | 13 | 44 | -11 |

### Supplement 6. Demographics of MDD patients, entire sample

|  | **Whole Patient Sample (N = 98)** | **CT Patient Sample (N = 74)** | **SSRI Patient Sample (N = 24)** |
| --- | --- | --- | --- |
| Gender | Female (N = 69, 70.41%) | Female (N = 52, 70.27%) | Female (N = 17, 70.83%) |
| Age | 35.60 (10.85) years | 35.32 (10.39) years | 36.46 (12.35) years |
| Ethnicity | Caucasian (N = 76, 77.55%) Non-Caucasian (N = 22, 22.45%) | Caucasian (N = 58, 78.38%) Non-Caucasian (N = 16, 21.62%) | Caucasian (N = 18, 75.00%) Non-Caucasian (N = 6, 25.00%) |
| Education | 14.98 (2.33) years | 15.36 (2.34) years* | 13.83 (1.90) years* |
| Depressive episodes | 3.54 (2.62) | 3.68 (2.66) | 3.14 (2.51) |
| Beck Depression Inventory - II (BDI-II) | Pre Treatment: 30.93 (9.04)  Post Treatment: 12.39 (9.74) | Pre Treatment: 31.48 (8.84)  Post Treatment: 12.73 (9.78) | Pre Treatment: 29.24 (9.62)  Post Treatment: 11.36 (9.75) |

Note. Means for age, education, number of depressive episodes, and Hamilton Depression Rating Scale are followed by standard deviations in parentheses. * Denotes that the CT and SSRI patient samples statistically differ on this variable. Number of depressive episodes above the 80^th^ percentile (8 episodes) of the entire sample were winsorized. CT sample education (N=72), number of depressive episodes (N=68), BDI-II Post (N=68). SSRI sample BDI post (N=22), number of depressive episodes (N=22). As in the primary sample, the CBT and SSRI samples differed by years of education, with the CBT sample reporting more years of education than the SSRI sample, t(94) = 2.89, p = .005, Hedges’ g = 0.68.

### Supplement 7. Application of meta-analytically derived regions to entire clinical dataset (not limited to those who completed treatment)

The sgACC ROI (derived from CBT studies) was prognostic of BDI change scores (*R*^2^ = .06, *F*(1,66) = 4.33, *p* = 0.041) in the whole CBT sample. Less reactivity was associated with greater improvement in symptoms. The relationship failed to reach statistical significance for residuals (*R*^2^ = .06, *F*(1,66) = 3.77, *p* = 0.057). This ROI did not predict BDI change scores (*R*^2^ <.01, *F*(1,21) = 0.01, *p* = 0.929) or residuals (*R*^2^ <.01, *F*(1,21) = 0.02, *p* = 0.902) for patients who received SSRIs. The pgACC ROI (derived from SSRI studies) did not significantly predict BDI change (*R*^2^ = 0.06, *F*(1,21) = 1.21, *p* = 0.284) or residuals (*R*^2^ = 0.09, *F*(1,21) = 2.08, *p* = 0.165) in the entire SSRI sample. Similar to the analyses among those who completed treatment, higher reactivity was non-significantly associated with higher residual symptomatology (in contrast to the literature). The ROI also did not predict treatment response for CBT patients, with negligible effect sizes for the entire sample (residuals: *R*^2^ = 0.01, *F*(1,67) = 0.44, *p* = 0.508; change scores: *R*^2^ < 0.01, *F*(1,67) = 0.17, *p* = 0.682).

### Supplement 8. Therapy ROI (subgenual cingulate) and BDI change scores for entire sample

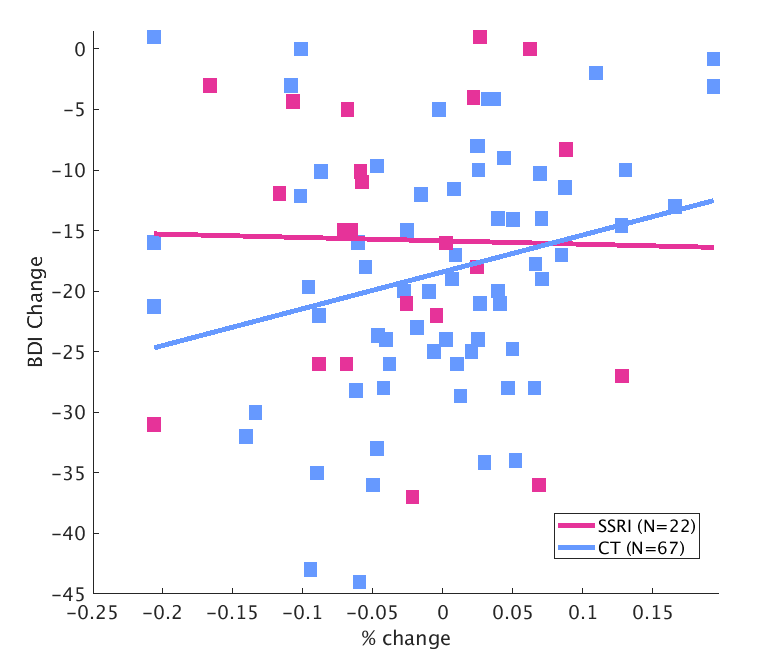

### Supplement 9. Prognostic accuracy of neural responses to faces for SSRI response in the empirical sample

**Introduction**

Results of the meta-analysis were potentially confounded by differences in stimulus modality between therapy studies, which generally used linguistic stimuli, and medication studies, which generally used pictorial stimuli, particularly, faces. Thus, as the rostral cingulate region’s activity to words was not prognostic for response to SSRIs in our primary analyses, we considered whether its responses to pictures might be prognostic. Largely by happenstance, we had collected neural reactivity to Hariri’s “Hammer” task (Hariri et al., 2000), which has reasonable psychometric properties in some regions (Sauder et al., 2013), on the same individuals, during the same fMRI scan, as words. Thus, this supplement considers the extent to which rostral cingulate responses to negative emotional faces on this task are associated with response in our SSRI sample.

**Methods**

**Participants.** The same participants who received SSRI treatment in the primary manuscript were examined.

**Task.** Hariri’s “Hammer” task (Hariri et al., 2000) was employed. Participants completed nine blocks in which they alternately matched a shape at the top of the screen to one of two shapes below it, in a triangle, or a face at the top of the screen to one of two faces below it. All three faces or shapes were shown simultaneously. Stimuli within blocks were randomly presented 1.5, 3, or 4.5 seconds apart, with shapes blocks containing 19 stimuli and faces blocks containing 25 stimuli. The block order was shapes, negative faces (fear or sad, from the Eckman stimulus set), shapes, neutral faces, shapes, negative faces, shapes, neutral faces, shapes. Thus, participants completed two blocks of negative faces.

**Analysis methods.** Data were pre-processed as described in the primary manuscript. Single subject analyses used canonical hemodynamic responses (AFNI’s “block” model) convolved with block onsets for each type of stimulus. Unstandardized beta weights were subjected to voxelwise outlier rescaling. Voxelwise regressions of the unstandardized beta weights (“neural responses”) to fearful faces, against residual Hamilton Depression Rating Scale and Beck Depression Inventory scores were performed. Type I error was protected for small volume correction within a mask comprising the pgACC in the VMPFC and the sgACC within BA25. Clusters significant at voxelwise p<.005 with an a priori cluster threshold at the maximum of the value empirically determined by AFNI’s 3dClustSim ACF option or 20 voxels (to account for the potential that 3dClustSim underestimates contiguity for small masks). 3dClustSim yielded an empirical region size of 7 voxels; thus a 20 voxel contiguity threshold was employed.

**Results**

Voxelwise regressions on residual Hamilton Rating Scale for Depression scores from neural reactivity to fearful faces in SSRI sample were predictive in both the pgACC (R2 = 0.28, F(1,18) = 6.67, p = 0.02) and sgACC (R2 = 0.32, F(1,18) = 8.13, p = 0.01), as shown in the figure below.

**Figure:** Relationship between pgACC and sgACC reactivity (regions derived from the whole-brain regressions of Supplement 9) and residual Hamilton Rating Scale for Depression scores in SSRI sample

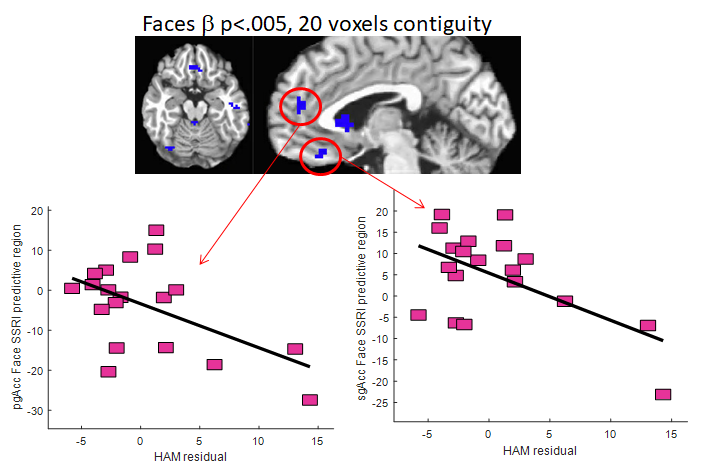

The observed pgAcc region did not overlap with the meta-analytically detected prognostic pgAcc region.

No significant subregions within the ACC were observed in relationship to residual Beck Depression Inventory - II scores.

**Discussion**

Consistent with the idea that prognostic accuracy may be a joint function of treatment medium and the employed task, whereas linguistic stimuli were not prognostic, for emotional faces, pre-treatment pgACC and sgACC cingulate regions were observed to be associated with residual depressive symptomatology.

### References to the Supplement

Alexopoulos, G. S., Hoptman, M. J., Kanellopoulos, D., Murphy, C. F., Lim, K. O., & Gunning, F. M. (2012). Functional connectivity in the cognitive control network and the default mode network in late-life depression. *Journal of Affective Disorders*, *139*(1), 56–65.

[Amsterdam, J. D., Newberg, A. B., Newman, C. F., Shults, J., Wintering, N., & Soeller, I. (2013). Change over time in brain serotonin transporter binding in major depression: effects of therapy measured with [123I]-ADAM SPECT. *Journal of Neuroimaging: Official Journal of the American Society of Neuroimaging*, *23*(4), 469–476.](http://paperpile.com/b/NFu4y0/e87aG)

Andreescu, C., Tudorascu, D. L., Butters, M. A., Tamburo, E., Patel, M., Price, J., Karp, J. F., Reynolds, C. F., 3rd, & Aizenstein, H. (2013). Resting state functional connectivity and treatment response in late-life depression. *Psychiatry Research*, *214*(3), 313–321.

Baldwin, R., Jeffries, S., Jackson, A., Sutcliffe, C., Thacker, N., Scott, M., & Burns, A. (2004). Treatment response in late-onset depression: relationship to neuropsychological, neuroradiological and vascular risk factors. *Psychological Medicine*, *34*(1), 125–136.

Beck, A. T., John Rush, A., Shaw, B. F., & Emery, G. (1979). *Cognitive Therapy of Depression*. Guilford Press.

Brannan, S. K., Mayberg, H. S., McGinnis, S., Silva, J. A., Tekell, J., Mahurin, R. K., Jerabek, P. A., & Fox, P. T. (2000). 355. Cingulate metabolism predicts treatment response: a replication. *Biological Psychiatry*, *47*(8), S107.

Brody, A. L., Saxena, S., Silverman, D. H., Alborzian, S., Fairbanks, L. A., Phelps, M. E., Huang, S. C., Wu, H. M., Maidment, K., & Baxter, L. R., Jr. (1999). Brain metabolic changes in major depressive disorder from pre- to post-treatment with paroxetine. *Psychiatry Research*, *91*(3), 127–139.

Carl, H., Walsh, E., Eisenlohr-Moul, T., Minkel, J., Crowther, A., Moore, T., Gibbs, D., Petty, C., Bizzell, J., Dichter, G. S., & Smoski, M. J. (2016). Sustained anterior cingulate cortex activation during reward processing predicts response to psychotherapy in major depressive disorder. *Journal of Affective Disorders*, *203*, 204–212.

Costafreda, S. G., Chu, C., Ashburner, J., & Fu, C. H. Y. (2009). Prognostic and diagnostic potential of the structural neuroanatomy of depression. *PloS One*, *4*(7). <https://www.ncbi.nlm.nih.gov/pmc/articles/pmc2712086/>

Crane, N. A., Jenkins, L. M., Bhaumik, R., Dion, C., Gowins, J. R., Mickey, B. J., Zubieta, J.-K., & Langenecker, S. A. (2017). Multidimensional prediction of treatment response to antidepressants with cognitive control and functional MRI. *Brain: A Journal of Neurology*, *140*(2), 472–486.

Cullen, K. R., Klimes-Dougan, B., Vu, D. P., Westlund Schreiner, M., Mueller, B. A., Eberly, L. E., Camchong, J., Westervelt, A., & Lim, K. O. (2016). Neural Correlates of Antidepressant Treatment Response in Adolescents with Major Depressive Disorder. *Journal of Child and Adolescent Psychopharmacology*, *26*(8), 705–712.

Davidson, R. J., Irwin, W., Anderle, M. J., & Kalin, N. H. (2003). The neural substrates of affective processing in depressed patients treated with venlafaxine. *The American Journal of Psychiatry*, *160*(1), 64–75.

Delaveau, P., Jabourian, M., Lemogne, C., Allaïli, N., Choucha, W., Girault, N., Lehericy, S., Laredo, J., & Fossati, P. (2016). Antidepressant short-term and long-term brain effects during self-referential processing in major depression. *Psychiatry Research. Neuroimaging*, *247*, 17–24.

Dunlop, B. W., Rajendra, J. K., Craighead, W. E., Kelley, M. E., McGrath, C. L., Choi, K. S., Kinkead, B., Nemeroff, C. B., & Mayberg, H. S. (2017). Functional Connectivity of the Subcallosal Cingulate Cortex And Differential Outcomes to Treatment With Cognitive-Behavioral Therapy or Antidepressant Medication for Major Depressive Disorder. *The American Journal of Psychiatry*, *174*(6), 533–545.

First, M. B., Spitzer, R. L., Gibbon, M., & Williams, J. B. W. (1995). Structured Clinical Interview for DSM-IV Axis I Disorders (SCID). New York, New York State Psychiatric Institute. *Biometrics Research*.

Frodl, T., Jäger, M., Smajstrlova, I., Born, C., Bottlender, R., Palladino, T., Reiser, M., Möller, H.-J., & Meisenzahl, E. M. (2008). Effect of hippocampal and amygdala volumes on clinical outcomes in major depression: a 3-year prospective magnetic resonance imaging study. *Journal of Psychiatry & Neuroscience: JPN*, *33*(5), 423–430.

Frodl, T., Meisenzahl, E. M., Zetzsche, T., Höhne, T., Banac, S., Schorr, C., Jäger, M., Leinsinger, G., Bottlender, R., Reiser, M., & Möller, H.-J. (2004). Hippocampal and amygdala changes in patients with major depressive disorder and healthy controls during a 1-year follow-up. *The Journal of Clinical Psychiatry*, *65*(4), 492–499.

Frodl, T., Scheuerecker, J., Schoepf, V., Linn, J., Koutsouleris, N., Bokde, A. L. W., Hampel, H., Möller, H.-J., Brückmann, H., Wiesmann, M., & Meisenzahl, E. (2011). Different effects of mirtazapine and venlafaxine on brain activation: an open randomized controlled fMRI study. *The Journal of Clinical Psychiatry*, *72*(4), 448–457.

Fu, C. H. Y., Costafreda, S. G., Sankar, A., Adams, T. M., Rasenick, M. M., Liu, P., Donati, R., Maglanoc, L. A., Horton, P., & Marangell, L. B. (2015). Multimodal functional and structural neuroimaging investigation of major depressive disorder following treatment with duloxetine. *BMC Psychiatry*, *15*, 82.

Furey, M. L., Drevets, W. C., Hoffman, E. M., Frankel, E., Speer, A. M., & Zarate, C. A., Jr. (2013). Potential of pretreatment neural activity in the visual cortex during emotional processing to predict treatment response to scopolamine in major depressive disorder. *JAMA Psychiatry* , *70*(3), 280–290.

Furey, M. L., Drevets, W. C., Szczepanik, J., Khanna, A., Nugent, A., & Zarate, C. A., Jr. (2015). Pretreatment Differences in BOLD Response to Emotional Faces Correlate with Antidepressant Response to Scopolamine. *The International Journal of Neuropsychopharmacology / Official Scientific Journal of the Collegium Internationale Neuropsychopharmacologicum* , *18*(8). https://doi.org/[10.1093/ijnp/pyv028](http://dx.doi.org/10.1093/ijnp/pyv028)

Godlewska, B. R., Browning, M., Norbury, R., Cowen, P. J., & Harmer, C. J. (2016). Early changes in emotional processing as a marker of clinical response to SSRI treatment in depression. *Translational Psychiatry*, *6*(11), e957.

Goldapple, K., Segal, Z., Garson, C., Lau, M., Bieling, P., Kennedy, S., & Mayberg, H. (2004). Modulation of cortical-limbic pathways in major depression: treatment-specific effects of cognitive behavior therapy. *Archives of General Psychiatry*, *61*(1), 34–41.

Gong, Q., Wu, Q., Scarpazza, C., Lui, S., Jia, Z., Marquand, A., Huang, X., McGuire, P., & Mechelli, A. (2011). Prognostic prediction of therapeutic response in depression using high-field MR imaging. *NeuroImage*, *55*(4), 1497–1503.

Gunning-Dixon, F. M., Walton, M., Cheng, J., Acuna, J., Klimstra, S., Zimmerman, M. E., Brickman, A. M., Hoptman, M. J., Young, R. C., & Alexopoulos, G. S. (2010). MRI signal hyperintensities and treatment remission of geriatric depression. *Journal of Affective Disorders*, *126*(3), 395–401.

Gyurak, A., Patenaude, B., Korgaonkar, M. S., Grieve, S. M., Williams, L. M., & Etkin, A. (2016). Frontoparietal Activation During Response Inhibition Predicts Remission to Antidepressants in Patients With Major Depression. *Biological Psychiatry*, *79*(4), 274–281.

Hamilton, M. (1960). A rating scale for depression. *Journal of Neurology, Neurosurgery, and Psychiatry*, *23*, 56–62.

Hariri, A. R., Bookheimer, S. Y., & Mazziotta, J. C. (2000). Modulating emotional responses: effects of a neocortical network on the limbic system. *Neuroreport*, *11*(1), 43–48.

Hernández-Ribas, R., Deus, J., Pujol, J., Segalàs, C., Vallejo, J., Menchón, J. M., Cardoner, N., & Soriano-Mas, C. (2013). Identifying brain imaging correlates of clinical response to repetitive transcranial magnetic stimulation (rTMS) in major depression. *Brain Stimulation*, *6*(1), 54–61.

He, Z., Cui, Q., Zheng, J., Duan, X., Pang, Y., Gao, Q., Han, S., Long, Z., Wang, Y., Li, J., Wang, X., Zhao, J., & Chen, H. (2016). Frequency-specific alterations in functional connectivity in treatment-resistant and -sensitive major depressive disorder. *Journal of Psychiatric Research*, *82*, 30–39.

[Hirvonen, J., Hietala, J., Kajander, J., Markkula, J., Rasi-Hakala, H., Salminen, J. K., Någren, K., Aalto, S., & Karlsson, H. (2011). Effects of antidepressant drug treatment and psychotherapy on striatal and thalamic dopamine D2/3 receptors in major depressive disorder studied with [11C] raclopride PET. *Journal of Psychopharmacology* , *25*(10), 1329–1336.](http://paperpile.com/b/NFu4y0/ZgS6H)

Hoogenboom, W. S., Perlis, R. H., Smoller, J. W., Zeng-Treitler, Q., Gainer, V. S., Murphy, S. N., Churchill, S. E., Kohane, I. S., Shenton, M. E., & Iosifescu, D. V. (2014). Limbic system white matter microstructure and long-term treatment outcome in major depressive disorder: a diffusion tensor imaging study using legacy data. *The World Journal of Biological Psychiatry: The Official Journal of the World Federation of Societies of Biological Psychiatry*, *15*(2), 122–134.

Hsieh, M.-H., McQuoid, D. R., Levy, R. M., Payne, M. E., MacFall, J. R., & Steffens, D. C. (2002). Hippocampal volume and antidepressant response in geriatric depression. *International Journal of Geriatric Psychiatry*, *17*(6), 519–525.

Iosifescu, D. V., Renshaw, P. F., Lyoo, I. K., Lee, H. K., Perlis, R. H., Papakostas, G. I., Nierenberg, A. A., & Fava, M. (2006). Brain white-matter hyperintensities and treatment outcome in major depressive disorder. *The British Journal of Psychiatry: The Journal of Mental Science*, *188*, 180–185.

Jarrett, R. B., Minhajuddin, A., Gershenfeld, H., Friedman, E. S., & Thase, M. E. (2013). Preventing depressive relapse and recurrence in higher-risk cognitive therapy responders: a randomized trial of continuation phase cognitive therapy, fluoxetine, or matched pill placebo. *JAMA Psychiatry* , *70*(11), 1152–1160.

Jung, J., Kang, J., Won, E., Nam, K., Lee, M.-S., Tae, W. S., & Ham, B.-J. (2014). Impact of lingual gyrus volume on antidepressant response and neurocognitive functions in Major Depressive Disorder: a voxel-based morphometry study. *Journal of Affective Disorders*, *169*, 179–187.

Keedwell, P. A., Drapier, D., Surguladze, S., Giampietro, V., Brammer, M., & Phillips, M. (2010). Subgenual cingulate and visual cortex responses to sad faces predict clinical outcome during antidepressant treatment for depression. *Journal of Affective Disorders*, *120*(1-3), 120–125.

Kennedy, S. H., Konarski, J. Z., Segal, Z. V., Lau, M. A., Bieling, P. J., McIntyre, R. S., & Mayberg, H. S. (2007). Differences in brain glucose metabolism between responders to CBT and venlafaxine in a 16-week randomized controlled trial. *The American Journal of Psychiatry*, *164*(5), 778–788.

Khalaf, A., Edelman, K., Tudorascu, D., Andreescu, C., Reynolds, C. F., & Aizenstein, H. (2015). White Matter Hyperintensity Accumulation During Treatment of Late-Life Depression. *Neuropsychopharmacology: Official Publication of the American College of Neuropsychopharmacology*, *40*(13), 3027–3035.

Konarski, J. Z., Kennedy, S. H., Segal, Z. V., Lau, M. A., Bieling, P. J., McIntyre, R. S., & Mayberg, H. S. (2009). Predictors of nonresponse to cognitive behavioural therapy or venlafaxine using glucose metabolism in major depressive disorder. *Journal of Psychiatry & Neuroscience: JPN*, *34*(3), 175.

Korb, A. S., Hunter, A. M., Cook, I. A., & Leuchter, A. F. (2009). Rostral anterior cingulate cortex theta current density and response to antidepressants and placebo in major depression. *Clinical Neurophysiology: Official Journal of the International Federation of Clinical Neurophysiology*, *120*(7), 1313–1319.

Langenecker, S. A., Kennedy, S. E., Guidotti, L. M., Briceno, E. M., Own, L. S., Hooven, T., Young, E. A., Akil, H., Noll, D. C., & Zubieta, J.-K. (2007). Frontal and limbic activation during inhibitory control predicts treatment response in major depressive disorder. *Biological Psychiatry*, *62*(11), 1272–1280.

Li, C.-T., Lin, C.-P., Chou, K.-H., Chen, I.-Y., Hsieh, J.-C., Wu, C.-L., Lin, W.-C., & Su, T.-P. (2010). Structural and cognitive deficits in remitting and non-remitting recurrent depression: a voxel-based morphometric study. *NeuroImage*, *50*(1), 347–356.

Lisiecka, D., Meisenzahl, E., Scheuerecker, J., Schoepf, V., Whitty, P., Chaney, A., Moeller, H.-J., Wiesmann, M., & Frodl, T. (2011). Neural correlates of treatment outcome in major depression. *The International Journal of Neuropsychopharmacology / Official Scientific Journal of the Collegium Internationale Neuropsychopharmacologicum* , *14*(4), 521–534.

Little, J. T., Ketter, T. A., Kimbrell, T. A., Dunn, R. T., Benson, B. E., Willis, M. W., Luckenbaugh, D. A., & Post, R. M. (2005). Bupropion and venlafaxine responders differ in pretreatment regional cerebral metabolism in unipolar depression. *Biological Psychiatry*, *57*(3), 220–228.

Liu, F., Guo, W., Yu, D., Gao, Q., Gao, K., Xue, Z., Du, H., Zhang, J., Tan, C., Liu, Z., Zhao, J., & Chen, H. (2012). Classification of different therapeutic responses of major depressive disorder with multivariate pattern analysis method based on structural MR scans. *PloS One*, *7*(7), e40968.

López-Solà, M., Pujol, J., Hernández-Ribas, R., Harrison, B. J., Contreras-Rodríguez, O., Soriano-Mas, C., Deus, J., Ortiz, H., Menchón, J. M., Vallejo, J., & Cardoner, N. (2010). Effects of duloxetine treatment on brain response to painful stimulation in major depressive disorder. *Neuropsychopharmacology: Official Publication of the American College of Neuropsychopharmacology*, *35*(11), 2305–2317.

Mackin, R. S., Tosun, D., Mueller, S. G., Lee, J.-Y., Insel, P., Schuff, N., Truran-Sacrey, D., Arean, P., Nelson, J. C., & Weiner, M. W. (2013). Patterns of reduced cortical thickness in late-life depression and relationship to psychotherapeutic response. *The American Journal of Geriatric Psychiatry: Official Journal of the American Association for Geriatric Psychiatry*, *21*(8), 794–802.

MacQueen, G. M., Yucel, K., Taylor, V. H., Macdonald, K., & Joffe, R. (2008). Posterior hippocampal volumes are associated with remission rates in patients with major depressive disorder. *Biological Psychiatry*, *64*(10), 880–883.

Marquand, A. F., Mourão-Miranda, J., Brammer, M. J., Cleare, A. J., & Fu, C. H. Y. (2008). Neuroanatomy of verbal working memory as a diagnostic biomarker for depression. *Neuroreport*, *19*(15), 1507–1511.

Mayberg, H. S., Brannan, S. K., Tekell, J. L., Silva, J. A., Mahurin, R. K., McGinnis, S., & Jerabek, P. A. (2000). Regional metabolic effects of fluoxetine in major depression: serial changes and relationship to clinical response. *Biological Psychiatry*, *48*(8), 830–843.

McGrath, C. L., Kelley, M. E., Dunlop, B. W., Holtzheimer, P. E., 3rd, Craighead, W. E., & Mayberg, H. S. (2014). Pretreatment brain states identify likely nonresponse to standard treatments for depression. *Biological Psychiatry*, *76*(7), 527–535.

McGrath, C. L., Kelley, M. E., Holtzheimer, P. E., Dunlop, B. W., Craighead, W. E., Franco, A. R., Craddock, R. C., & Mayberg, H. S. (2013). Toward a neuroimaging treatment selection biomarker for major depressive disorder. *JAMA Psychiatry* , *70*(8), 821–829.

Milak, M. S., Parsey, R. V., Lee, L., Oquendo, M. A., Olvet, D. M., Eipper, F., Malone, K., & Mann, J. J. (2009). Pretreatment regional brain glucose uptake in the midbrain on PET may predict remission from a major depressive episode after three months of treatment. *Psychiatry Research*, *173*(1), 63–70.

Mulert, C., Juckel, G., Brunnmeier, M., Karch, S., Leicht, G., Mergl, R., Möller, H.-J., Hegerl, U., & Pogarell, O. (2007). Prediction of treatment response in major depression: integration of concepts. *Journal of Affective Disorders*, *98*(3), 215–225.

Nolen-Hoeksema, S. (1991). Responses to depression and their effects on the duration of depressive episodes. *Journal of Abnormal Psychology*, *100*(4), 569–582.

Patankar, T. F., Baldwin, R., Mitra, D., Jeffries, S., Sutcliffe, C., Burns, A., & Jackson, A. (2007). Virchow–Robin space dilatation may predict resistance to antidepressant monotherapy in elderly patients with depression. *Journal of Affective Disorders*, *97*(1), 265–270.

Phillips, J. L., Batten, L. A., Aldosary, F., Tremblay, P., & Blier, P. (2012). Brain-volume increase with sustained remission in patients with treatment-resistant unipolar depression. *The Journal of Clinical Psychiatry*, *73*(5), 625–631.

Pizzagalli, D., Pascual-Marqui, R. D., Nitschke, J. B., Oakes, T. R., Larson, C. L., Abercrombie, H. C., Schaefer, S. M., Koger, J. V., Benca, R. M., & Davidson, R. J. (2001). Anterior cingulate activity as a predictor of degree of treatment response in major depression: evidence from brain electrical tomography analysis. *The American Journal of Psychiatry*, *158*(3), 405–415.

Rizvi, S. J., Salomons, T. V., Konarski, J. Z., Downar, J., Giacobbe, P., McIntyre, R. S., & Kennedy, S. H. (2013). Neural response to emotional stimuli associated with successful antidepressant treatment and behavioral activation. *Journal of Affective Disorders*, *151*(2), 573–581.

Roffman, J. L., Witte, J. M., Tanner, A. S., Ghaznavi, S., Abernethy, R. S., Crain, L. D., Giulino, P. U., Lable, I., Levy, R. A., Dougherty, D. D., Evans, K. C., & Fava, M. (2014). Neural predictors of successful brief psychodynamic psychotherapy for persistent depression. *Psychotherapy and Psychosomatics*, *83*(6), 364–370.

Salvadore, G., Cornwell, B. R., Colon-Rosario, V., Coppola, R., Grillon, C., Zarate, C. A., & Manji, H. K. (2009). Increased Anterior Cingulate Cortical Activity in Response to Fearful Faces: A Neurophysiological Biomarker that Predicts Rapid Antidepressant Response to Ketamine. *Biological Psychiatry*, *65*(4), 289–295.

Samson, A. C., Meisenzahl, E., Scheuerecker, J., Rose, E., Schoepf, V., Wiesmann, M., & Frodl, T. (2011). Brain activation predicts treatment improvement in patients with major depressive disorder. *Journal of Psychiatric Research*, *45*(9), 1214–1222.

Sanacora, G., Fenton, L. R., Fasula, M. K., Rothman, D. L., Levin, Y., Krystal, J. H., & Mason, G. F. (2006). Cortical γ-Aminobutyric Acid Concentrations in Depressed Patients Receiving Cognitive Behavioral Therapy. *Biological Psychiatry*, *59*(3), 284–286.

Sauder, C. L., Hajcak, G., Angstadt, M., & Phan, K. L. (2013). Test-retest reliability of amygdala response to emotional faces. *Psychophysiology*, *50*(11), 1147–1156.

Saxena, S., Brody, A. L., Ho, M. L., Zohrabi, N., Maidment, K. M., & Baxter, L. R., Jr. (2003). Differential brain metabolic predictors of response to paroxetine in obsessive-compulsive disorder versus major depression. *The American Journal of Psychiatry*, *160*(3), 522–532.

Siegle, G. J., Carter, C. S., & Thase, M. E. (2006). Use of FMRI to predict recovery from unipolar depression with cognitive behavior therapy. *The American Journal of Psychiatry*, *163*(4), 735–738.

Siegle, G. J., Thompson, W. K., Collier, A., Berman, S. R., Feldmiller, J., Thase, M. E., & Friedman, E. S. (2012). Toward clinically useful neuroimaging in depression treatment: prognostic utility of subgenual cingulate activity for determining depression outcome in cognitive therapy across studies, scanners, and patient characteristics. *Archives of General Psychiatry*, *69*(9), 913–924.

Spielberger, C. D. (2010). State‐Trait anxiety inventory. *The Corsini Encyclopedia of Psychology*. <https://onlinelibrary.wiley.com/doi/abs/10.1002/9780470479216.corpsy0943>

Straub, J., Metzger, C. D., Plener, P. L., Koelch, M. G., Groen, G., & Abler, B. (2017). Successful group psychotherapy of depression in adolescents alters fronto-limbic resting-state connectivity. *Journal of Affective Disorders*, *209*, 135–139.

Szczepanik, J., Nugent, A. C., Drevets, W. C., Khanna, A., Zarate, C. A., Jr, & Furey, M. L. (2016). Amygdala response to explicit sad face stimuli at baseline predicts antidepressant treatment response to scopolamine in major depressive disorder. *Psychiatry Research. Neuroimaging*, *254*, 67–73.

Thompson, D. G., Kesler, S. R., Sudheimer, K., Mehta, K. M., Thompson, L. W., Marquett, R. M., Holland, J. M., Reiser, R., Rasgon, N., Schatzberg, A., & O’Hara, R. M. (2015). FMRI activation during executive function predicts response to cognitive behavioral therapy in older, depressed adults. *The American Journal of Geriatric Psychiatry: Official Journal of the American Association for Geriatric Psychiatry*, *23*(1), 13–22.

Tiger, M., Rück, C., Forsberg, A., Varrone, A., Lindefors, N., Halldin, C., Farde, L., & Lundberg, J. (2014). Reduced 5-HT1B receptor binding in the dorsal brain stem after cognitive behavioural therapy of major depressive disorder. *Psychiatry Research: Neuroimaging*, *223*(2), 164–170.

Toki, S., Okamoto, Y., Onoda, K., Matsumoto, T., Yoshimura, S., Kunisato, Y., Okada, G., Shishida, K., Kobayakawa, M., Fukumoto, T., Machino, A., Inagaki, M., & Yamawaki, S. (2014). Hippocampal activation during associative encoding of word pairs and its relation to symptomatic improvement in depression: A functional and volumetric MRI study. *Journal of Affective Disorders*, *152-154*, 462–467.

Vakili, K., Pillay, S. S., Lafer, B., Fava, M., Renshaw, P. F., Bonello-Cintron, C. M., & Yurgelun-Todd, D. A. (2000). Hippocampal volume in primary unipolar major depression: a magnetic resonance imaging study. *Biological Psychiatry*, *47*(12), 1087–1090.

Wagner, G., Koch, K., Schachtzabel, C., Sobanski, T., Reichenbach, J. R., Sauer, H., & Schlösser, R. G. M. (2010). Differential effects of serotonergic and noradrenergic antidepressants on brain activity during a cognitive control task and neurofunctional prediction of treatment outcome in patients with depression. *Journal of Psychiatry & Neuroscience: JPN*, *35*(4), 247–257.

Walsh, N. D., Williams, S. C. R., Brammer, M. J., Bullmore, E. T., Kim, J., Suckling, J., Mitterschiffthaler, M. T., Cleare, A. J., Pich, E. M., Mehta, M. A., & Fu, C. H. Y. (2007). A longitudinal functional magnetic resonance imaging study of verbal working memory in depression after antidepressant therapy. *Biological Psychiatry*, *62*(11), 1236–1243.

Wang, L.-J., Kuang, W.-H., Xu, J.-J., Lei, D., & Yang, Y.-C. (2014). Resting-state brain activation correlates with short-time antidepressant treatment outcome in drug-naive patients with major depressive disorder. *The Journal of International Medical Research*, *42*(4), 966–975.

Wang, L., Xia, M., Li, K., Zeng, Y., Su, Y., Dai, W., Zhang, Q., Jin, Z., Mitchell, P. B., Yu, X., & Others. (2015). The effects of antidepressant treatment on resting-state functional brain networks in patients with major depressive disorder. *Human Brain Mapping*, *36*(2), 768–778.

Watson, D., Weber, K., Assenheimer, J. S., Clark, L. A., Strauss, M. E., & McCormick, R. A. (1995). Testing a tripartite model: I. Evaluating the convergent and discriminant validity of anxiety and depression symptom scales. *Journal of Abnormal Psychology*, *104*(1), 3–14.

Williams, L. M., Korgaonkar, M. S., Song, Y. C., Paton, R., Eagles, S., Goldstein-Piekarski, A., Grieve, S. M., Harris, A. W. F., Usherwood, T., & Etkin, A. (2015). Amygdala Reactivity to Emotional Faces in the Prediction of General and Medication-Specific Responses to Antidepressant Treatment in the Randomized iSPOT-D Trial. *Neuropsychopharmacology: Official Publication of the American College of Neuropsychopharmacology*, *40*(10), 2398–2408.

Yeh, Y. W., Ho, P. S., Kuo, S. C., & Chen, C. Y. (2015). Disproportionate reduction of serotonin transporter may predict the response and adherence to antidepressants in patients with major depressive disorder: a positron …. *Aquatic Microbial Ecology: International Journal*. <https://academic.oup.com/ijnp/article-abstract/18/7/pyu120/675478>
